# Attenuated conditioned taste aversion for sucrose in female mice with a history of chronic low-dose ethanol exposure

**DOI:** 10.64898/2026.06.08.730505

**Authors:** Christina M Curran-Alfaro, Christine M Side, Ankith Alluri, Wesley Corey, Conor Sheehan, Jacqueline M Barker

## Abstract

It is becoming increasingly clear that chronic exposure to lower levels of ethanol impact learning and behavior. To determine the impact of chronic low-dose ethanol exposure on sensitivity to changes in stimulus value, a conditioned taste aversion procedure was used. Adult male and female mice underwent a sucrose two bottle-choice drinking paradigm. Each day, mice received an injection of either low-dose ethanol (0.5g/kg) or saline two hours after sucrose access for 20 days. This was followed by a lithium chloride (LiCl)-induced conditioned taste aversion (CTA) paradigm in which 0.15M LiCl or vehicle injection was administered immediately after sucrose consumption for three days. On the fourth day, changes in sucrose consumption were analyzed. Chronic exposure to low-dose ethanol did not affect sucrose consumption in either female of male mice during two-bottle choice. In female mice, a history of chronic low-dose ethanol exposure blocked the development of LiCl-induced CTA. A history of chronic low-dose ethanol did not impact LiCl-induced CTA in male mice as both ethanol-naïve and -exposed male mice who underwent LiCl pairing reduced sucrose consumption. This suggests that low-dose ethanol alters aversion-related learning in female mice which may have implication for development of aberrant behavior and risk for alcohol use disorder (AUD).

## Introduction

The ability to update behavior in response to change is essential to flexibly navigate a changing environment (Lea et. al., 2020, Simoes et. al, 2012). Learning to avoid foods that cause aversive outcomes is common across species (Lin et al., 2017; Riley et al., 1985). This conditioned taste aversion (CTA) is a form of Pavlovian learning that requires individuals to associate post-ingestive outcomes – an unconditioned stimulus (US) - with the taste of the food consumed – the conditioned stimulus (CS). Reduced consumption is expected in CTA, which is thought to reflect reduced value of the taste of the CS. CTA thus requires learning of this new Pavlovian association to successfully devalue the taste (Rilet et al., 985; Adams & Dickinson, 1981).

Alcohol use disorder (AUD) is characterized by persistent drinking despite adverse consequences, and sensitivity to aversive experience can constrain alcohol use (King et al., 2011, 2019; Koob et al., 1997; Yang et al., 2022). Deficits in aversion learning may contribute to maintenance of ethanol seeking or consumption despite the aversive properties associated with ethanol exposure (King et al., 2011, 2019). Impaired expression or sensitivity to CTA has been identified as a potential deficit associated with increased risk for AUD (King et al., 2019; Przybysz, Ramirez, et al., 2024; Taxier et al., 2025). Chronic exposure to high doses of ethanol alters CTA in rodents, potentially due to altered aversion encoding or impaired acquisition of the new association (Glover et al., 2016; Lopez et al., 2014; Przybysz, Ramirez, et al., 2024; Przybysz, Shillinglaw, et al., 2024). This includes both suppressed incubation of CTA for nondrug reward (Ramirez et al., 2024) and resistance to lithium chloride devaluation of ethanol reward (Lopez et al., 2014) following chronic intermittent ethanol exposure – a model of ethanol dependence.

While a majority of research in this domain has emphasized the impact of higher doses of ethanol, there is increasing evidence that chronic exposure to lower doses of ethanol impact the brain and behavior, including learning and memory and reward-motivated behaviors (Bryant et al., 2022, 2023, 2024; Cui et al., 2017; Curran-Alfaro et al., 2026; Davidson et al., 1997; Daviet et al., 2022; Weafer et al., 2016). Previous work from our group has shown that chronic exposure to low-dose ethanol increased sensitivity to changes in reward magnitude in female mice in a task assessing motivation, suggesting that a history of lower dose ethanol exposure may be impacting sensitivity to changes in reward value. However, in the previous study, the reward value was manipulated within-session and did not require aversion learning, which is known to be impacted by chronic alcohol. To test the hypothesis that a history of chronic low dose ethanol exposure suppressed aversion learning, the current study utilized a lithium chloride (LiCl)-CTA paradigm, commonly used to assess learned devaluation. This work will further our understanding of how chronic exposure to low-dose ethanol alters reward-related behavior, including sensitivity to changes in reward value and aversion learning. This may suggest potential factors through which chronic exposure to lower doses of ethanol alters subsequent aversion sensitivity, which could ultimately confer risk for AUD or related outcomes.

## Results

### Chronic low-dose ethanol exposure does not impact 10% sucrose two-bottle choice consumption

To explore the effects of chronic low-dose ethanol exposure on 10% sucrose preference during a 20-day two-bottle choice paradigm (see behavioral timeline, **Fig 1A**), daily sucrose consumption was analyzed across days using two-way rmANOVA. In female mice, there was no impact of ethanol treatment on sucrose consumption [EtOH Condition: F (1, 22) = 0.2383, p = 0.6303; Days x EtOH Condition: F (4.530, 99.42) = 0.7085, p = 0.6049; Greenhouse-Geisser corrected, **Fig 1B**]. There was a main effect of day [F (4.530, 99.42) = 6.872, p <0.0001], but post hoc analysis did not reveal any significant differences in consumption vs day 1 [all p’s > 0.05, Sidak’s]. Mice were matched using sucrose consumption measures on day 20 for assignment to treatments groups for conditioned taste aversion (CTA). Analysis confirmed successful matching as consumption in the mice assigned to undergo LiCl treatment did not differ from controls [two-way ANOVA: *Females*: EtOH Condition: F (1, 20) = 0.0018, p = 0.9661; LiCl Treatment: F (1, 20) = 0.0002422, p = 0.9877; EtOH Condition x LiCl Treatment: F (1, 20) = 0.03783, p = 0.8477, **Fig 1C**].

**Figure 1.**
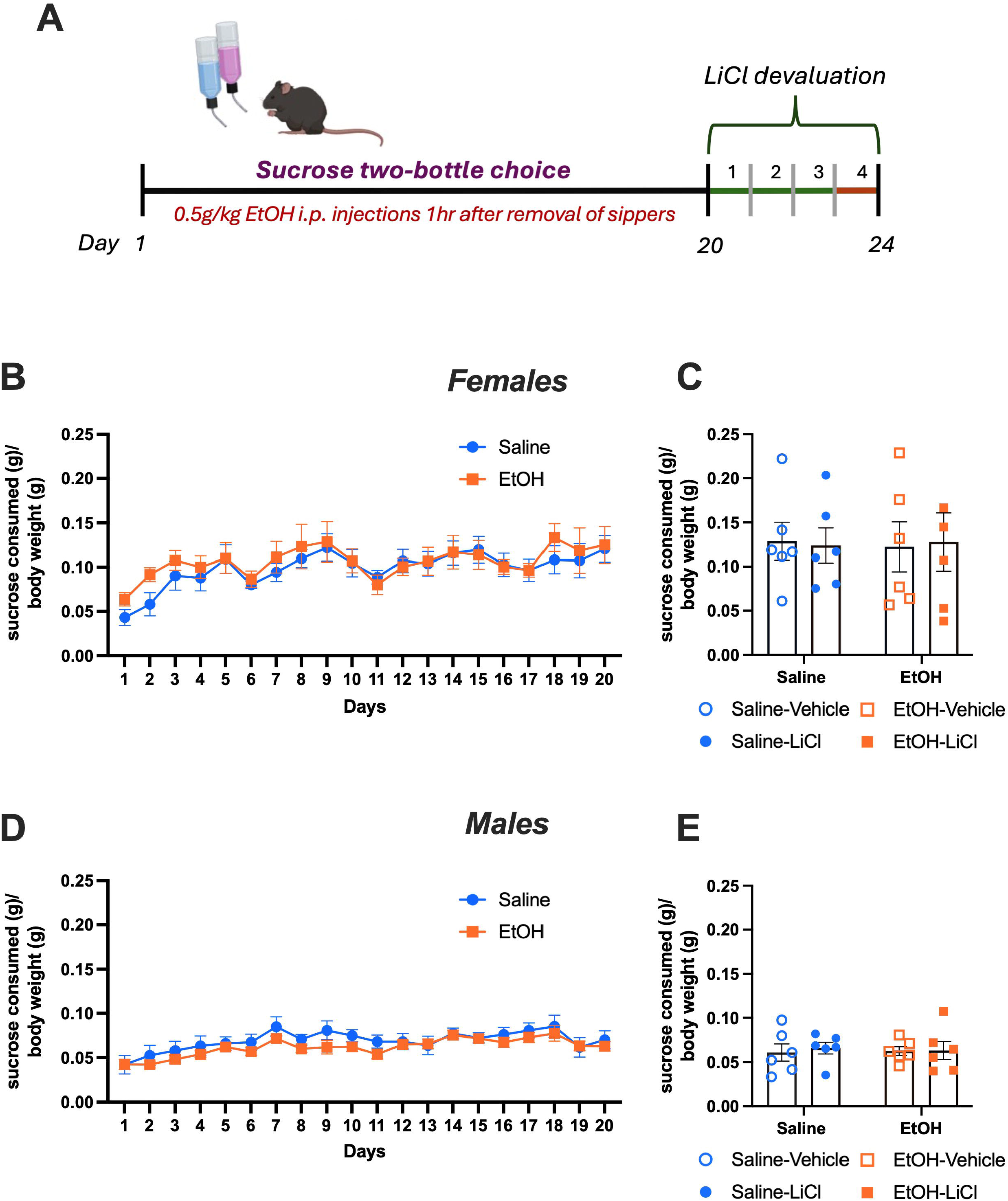
Chronic exposure to low-dose ethanol did not impact consumption in a sucrose two-bottle choice task. (**A**) Experimental behavioral timeline. (**B**) Chronic low-dose ethanol exposure did not impact intake in a sucrose two-bottle choice paradigm female mice. (**C**) Female mice were successfully matched by intake during the two-bottle choice paradigm to receive subsequent LiCl-CTA or serve as controls. (**D**) Chronic low-dose ethanol exposure did not impact intake in a sucrose two-bottle choice paradigm in male mice. (**E**) Male mice assigned to the LiCl-CTA and control groups had matched intake prior to LiCl conditioning. Females- saline: n = 12, ethanol: n = 12; males- saline: n = 12, ethanol: n = 12. Data represent mean +/- SEM.

Findings in male mice were similar to those observed in females. There was no effect of chronic low-dose ethanol on sucrose consumption across days [EtOH Condition: F (1, 22) = 0.8200, p = 0.3750; Days x Condition: F (6.241, 137.3) = 0.6314, p = 0.7112; Greenhouse-Geisser corrected, **Fig 1D**]. These was a main effect of day in male mice [F (6.241, 137.3) = 7.810, p <0.0001], as observed in female mice, yet post hoc analysis did not reveal any significant differences vs day 1 [all p’s > 0.05, Sidak’s]. Male mice were then matched using sucrose consumption on the final day of two-bottle choice to assignment treatment groups prior to the start of CTA treatment [EtOH Condition: F (1, 20) = 0.003253, p = 0.551; LiCl Treatment: F (1, 20) = 0.1186, p = 0.7342; EtOH Condition x LiCl Treatment: F (1, 20) = 0.06899, p = 0.7955, **Fig 1E**].

### Chronic low-dose ethanol impaired conditioned taste aversion expression in female mice

To investigate the effects of a history of chronic low-dose ethanol on CTA, mice underwent a 3-day pairing of LiCl treatment following an hour of 10% sucrose exposure [behavioral timeline, **Fig 2A**]. Sucrose consumption was analyzed following 3 days of LiCl treatment [**Fig 2B**]. In female mice, an interaction of chronic low-dose ethanol exposure and LiCl treatment was observed [two-way ANOVA: EtOH Condition: F (1, 20) = 0.5806, p = 0.4550; LiCl Treatment: F (1, 20) = 9.290, p = 0.0063; EtOH x LiCl Treatment: F (1, 20) = 6.452, p = 0.0195; **Fig 2C**]. In female mice *without* a history of low-dose ethanol exposure, mice that received LiCl pairing consumed less than controls [p = 0.0008], consistent with a CTA. However, in female mice with history of low-dose ethanol, there was no significant difference in consumption between mice who received LiCl pairing and controls [p = 0.7232], indicating the absence of a CTA. In contrast, this effect was not observed in male mice [**Fig 2D**]. Rather, male mice that had LiCl-paired with sucrose consumed less sucrose than saline-paired controls independent of low-dose ethanol exposure history [EtOH Condition: F (1, 20) = 0.2787, p = 0.6033; LiCl Treatment: F (1, 20) = 11.78, p = 0.0026; EtOH Condition x LiCl Treatment: F (1, 20) = 0.2787, p = 0.6033; **Fig 2E**], suggesting that male mice developed a CTA regardless of ethanol exposure history.

**Figure 2.**
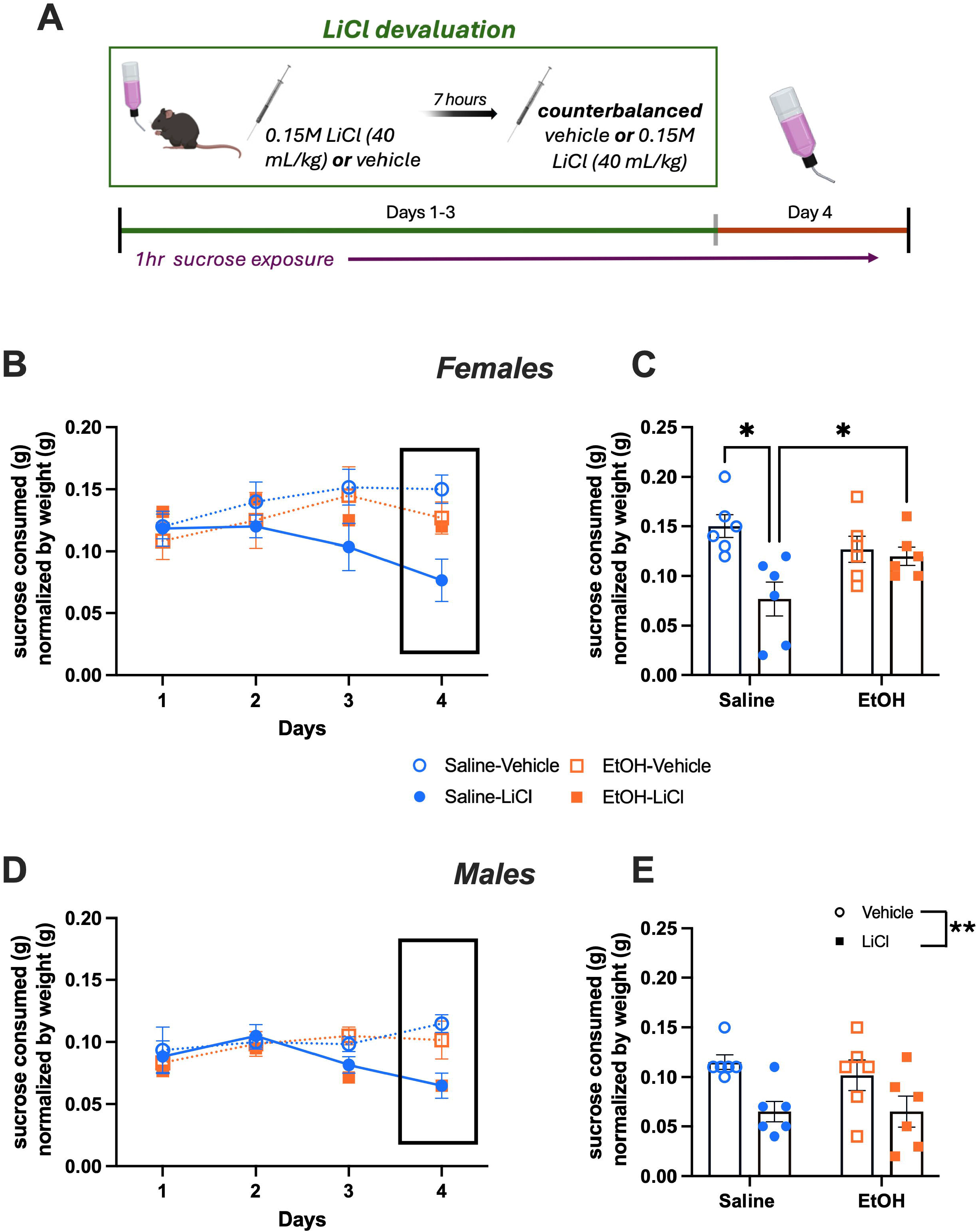
A history of chronic low-dose ethanol exposure blocked LiCl-CTA expression in female, not male, mice. (**A**) Timeline for conditioned LiCl devaluation. (**B**) A history of ethanol exposure interacted with LiCl pairing to alter sucrose consumption across LiCl-CTA, with ethanol-naive females exhibiting a reduction in sucrose intake compared to their controls. (**C**) On the final test day, ethanol-naïve females that underwent LiCl-CTA consumed less sucrose than all other groups. Low-dose ethanol-exposed females that underwent LiCl-CTA consumed similar amounts of sucrose as those that did not undergo LiCl-CTA. (**D**) A history of chronic low-dose ethanol did not affect expression of LiCl-CTA in male mice. (**E**) A main effect of LiCl paring was observed in male mice, regardless of ethanol exposure history. Females- saline-vehicle n = 6, saline-LiCl pairing n = 6, ethanol-vehicle: n = 6, ethanol-LiCl: n = 6; males- saline-vehicle n = 6, saline-LiCl pairing n = 6, ethanol-vehicle: n = 6, ethanol-LiCl. Data represent mean +/- SEM.

## Discussion

Even at low-doses, chronic ethanol exposure alters behavior, including promoting maladaptive motivated behavior and driving sensitivity to changes in reward magnitude, though this may occur in a sex-specific manner (Bryant et al., 2022; Curran-Alfaro et al., 2026). The current work investigated the effects of chronic, low-dose ethanol on conditioned sucrose reward devaluation. No effect of chronic low-dose ethanol was observed in either female or male mice during two-bottle choice consumption sessions prior to LiCl pairing, suggesting that low-dose ethanol does not impact sucrose preference or normal intake. While repeated low-dose ethanol did not impact baseline sucrose consumption, chronic exposure to low-dose ethanol blocked the expression of LiCl conditioned taste aversion (CTA) in female mice. This effect was not observed in male mice. The reduced expression of LiCl in female mice exposed to chronic low-dose ethanol may result from a failure to learn the LiCl-sucrose association and/or an impaired ability to update reward value, with implications for aversion learning.

Previous work has identified disrupted conditioned taste aversion following chronic exposure to higher doses of ethanol and predominantly been performed in males. Work utilizing chronic intermittent ethanol exposure (CIE), a preclinical model of ethanol dependence and withdrawal, has shown that male mice with a history of high dose ethanol did not express LiCl-CTA in a sweetened ethanol self-administration task (Lopez et al., 2014). It has also been demonstrated that CIE attenuated incubation of ethanol-CTA in comparison to ethanol-CTA expression prior to chronic high dose ethanol exposure suggesting that chronic exposure to higher doses of ethanol may impair the ability to update aversive associations, as opposed to attenuating the aversive properties of ethanol exposure itself (Ramirez et al., 2024). In contrast to dependence models, male mice who self-administered ethanol were sensitive to LiCl CTA, even with protracted self-administration timelines (10 week). This is in line with the current results, in which male mice exposed to low-dose ethanol for 20 days exhibited intact LiCl-CTA, suggesting that both self- and experimenter-administered low-dose ethanol exposure is not sufficient to alter LiCl-CTA in males.

Aligned with the current results, our previous work has shown increased sensitivity to changes in reward magnitude in female mice with a history of chronic low-dose ethanol exposure. Together, this suggests that female mice may be vulnerable to the effects of low-dose ethanol on reward value-driven behavior, while male mice which may require repeated exposure to higher doses to impact reward-value sensitivity. Notably, the previous findings demonstrated *increased* sensitivity to change in reward magnitude in female mice, while this work indicates resistance to LiCl devaluation. One difference may reflect the learned nature of the CTA versus the within-session manipulation of reward magnitude in the previous study (Curran-Alfaro et al., 2026). Here, the pairing of LiCl with sucrose required learning across conditioning days to update the sucrose association with the aversive experience. This may suggest that chronic low-dose ethanol exposure can alter reward revaluation in female mice. It is also possible that chronic low-dose ethanol exposure in female mice promotes an initial over-valuation of the sucrose reward, which results in an apparent increase sensitivity to changes in reward magnitude or resistance to aversion conditioning and learning. Alternatively, this may reflect sex differences in aversion sensitivity. For example, female mice have been shown to have greater resistance to quinine adulteration of ethanol (or lowered sensitivity to the aversive tastant) compared to male mice, while female rats exhibit less aversive behavior in response to neophobia-inducing stimuli. Pairings of the aversive physical effects of ethanol, such as motor impairment and intoxication with food consumption can induce a CTA (Przybysz, Ramirez, et al., 2024; Rabin et al., 1989; Ramirez et al., 2024; Taxier et al., 2025). In both male and female rats, ethanol produced similar CTA expression to that of LiCl. Acute exposure to ethanol prior to ethanol-CTA attenuated ethanol-CTA, but not LiCl-CTA, in males suggesting that repeated ethanol exposure may be necessary to impact LiCl-CTA (Rabin et al., 1989). However, these studies did not explore the effects of ethanol on LiCl-CTA expression in females.

Ethanol-induced CTA requires common neural substrates to LiCl-CTA, and it is possible that repeated low-dose ethanol exposure may disrupt normal activity within these substrates, particularly in female mice who exhibited reduced LiCl-CTA after exposure. These targets include the rostromedial tegmental nucleus (RMTg), ventral tegmental area (VTA), and the insular cortex (IC). Expression of cFos within the RMTg is increased following LiCl-CTA and ethanol-induced CTA in both female and male rats (Glover et al., 2016). Lesions to the RMTg accelerate extinction of ethanol-induced CTA, resulting in an increase of ethanol consumption (Sheth et al., 2016). The RMTg is also considered a “master brake” of the ventral tegmental area (VTA), a contributor to aversive learning. Female mice exhibit greater VTA activity following consumption of quinine-adulterated ethanol, which may suggest less sensitivity of this circuit to aversive tastants, leading to reduced suppression of consumption of quinine-adulterated ethanol in females in comparison to male mice (Arnold et al., 2023). Similarly, sex differences in activation of the IC, a structure necessary for both ethanol-induced and LiCl-CTA, have been reported (Kayyal et al., 2019). For example, female mice displayed a decrease in activity within the posterior IC during aversion learning compared to male mice, while altering GABAergic tone within the anterior IC reduced CTA strength in male, but not female, mice. These regulators of CTA are known to be sensitive to the effects of higher doses of ethanol exposure (Chen et al., 2015; Glover et al., 2019; Hopf et al., 2007; Mitten et al., 2025; Morikawa et al., 2010; Mukherjee et al., 2023; Przybysz, Shillinglaw, et al., 2024), however, the impact of chronic low-dose ethanol on these substrates is not well understood.

### Caveats and future considerations

The current work demonstrated behavioral impacts of chronic low-dose ethanol exposure on sucrose LiCl-CTA expression in female mice. One caveat is that ethanol exposure was a bolus injection administered by the experimenter. This method was selected to control for dose and to purposefully delay timing of ethanol exposure during sucrose two-bottle choice to avoid an ethanol-induced CTA. Findings suggest that this was avoided as ethanol-treated mice consumed similar sucrose to saline controls prior to LiCl CTA procedures, however it is possible that ethanol administration impacted sucrose value and this cannot be ruled out in the current findings. While intraperitoneal injection also minimized stress compared to oral gavage, it is possible that the post-ingestive effects of ethanol administered orally would yield different outcomes than intraperitoneal injection. While the current experiments identify behavioral alterations, future work should explore the underlying neural mechanisms by which chronic low-dose ethanol impacts ethanol-induced CTA. The addition of mechanistic approaches would be useful to further characterize the distinct effects of chronic low-dose versus higher-dose ethanol on brain and aversion learning. Additional behavioral studies would be beneficial to identify how chronic low-dose ethanol exposure in female mice may be impairing underlying processes necessary for LiCl-CTA expression. Importantly, this study cannot dissociate chronic low-dose ethanol effects on sensitivity to LiCl-induced aversion versus impaired aversion learning, which merits future consideration. Finally, these results should be expanded to determine how chronic low-dose ethanol exposure impacts the development of ethanol-induced CTA to determine whether these effects generalize to other aversive learning models.

### Conclusions

These findings build on a growing literature identifying consequences of chronic low-dose ethanol exposure on reward-related behavior. In this study, chronic exposure to low-dose ethanol induced sex-specific effects, with ethanol-exposed female mice exhibiting blunted LiCl-CTA expression. This work highlights the importance of further understanding the unique effects of chronic low-dose ethanol and sex on reward-related behaviors, such as aversion sensitivity or learning, to characterize phenotypes which may be at a greater risk for alcohol-associated negative outcomes.

## Methods and Materials

### Subjects

Adult C57BL/6J mice from Jackson Laboratory (delivered at 9 weeks in age; *N* = 48; 24 females, 24 males) were used in the following studies. Studies were performed in accordance with protocols approved by the Drexel University Institutional Animal Care and Use Committee. Mice were housed in a standard 12h:12h light-dark cycle vivarium. Mice were allowed to acclimate to the facilities for at least one week and housed individually before the start of the experiments.

### Sucrose two-bottle choice

Mice underwent a 10% sucrose two-bottle choice paradigm in which two glass 10mL sipper bottles (Amuza Inc, #H37.100.01), one containing water and the other with 10% sucrose, were placed in each animal’s home cage. Both water and sucrose sipper bottles were weighed prior to being placed within each respective home cage for 2 hours for 20 days, approximately 2 hours into the light cycle. Bottles were counterbalanced each day to alternating sides and were weighed immediately upon removal to record consumption.

### Low-dose ethanol exposure

Mice were matched by sex to receive low-dose ethanol (0.5g/kg, i.p.) or saline vehicle injections 1 hour after removal of sipper bottles for all 20 days of 10% sucrose consumption. After 20 days of 10% sucrose two-bottle choice was completed, animals no longer received ethanol or saline injections. Thus, all low-dose ethanol exposure was completed prior to beginning lithium chloride condition taste aversion conditioning.

### Conditioned Taste Aversion

On three consecutive days, mice underwent an hour of 10% sucrose exposure, approximately 2 hours into the light cycle, and then received either 0.15 LiCl (40mL/kg) or vehicle injections (matched by sex and condition) immediately upon removal of the 10% sucrose glass sipper bottle. Bottles were weighed for consumption. Seven hours later, counterbalanced injections were administered such that mice that received saline immediately after sucrose consumption received an injection of LiCl, and vice versa, to account for any off-targets effects of LiCl exposure.

### Statistical analysis

All statistical analyses were conducted using GraphPad Prism (Version 10.6.0). Data were analyzed using two-way ANOVA or, for repeated behavioral testing, rmANOVA. Given known sex differences in ethanol intake (Li et al., 2019; Middaugh et al., 1999; Vetter-O’Hagen et al., 2009), analyses were separated by sex to determine the effect of low-dose ethanol on intake within each sex, rather than sex differences in magnitude of effect.

## References

Adams, C. D., & Dickinson, A. (1981). Instrumental Responding following Reinforcer Devaluation. The Quarterly Journal of Experimental Psychology Section B, 33(2b), 109–121. 10.1080/14640748108400816

Arnold, M. E., Butts, A. N., Erlenbach, T. R., Amico, K. N., & Schank, J. R. (2023). Sex differences in neuronal activation during aversion-resistant alcohol consumption. Alcohol: Clinical and Experimental Research, 47(2), 240–250. doi: 10.1111/acer.15006

Bryant, K. G., Nothem, M. A., Buck, L. A., Singh, B., Amin, S., Curran-Alfaro, C. M., & Barker, J. M. (2023). A History of Low-Dose Ethanol Shifts the Role of Ventral Hippocampus during Reward Seeking in Male Mice. ENeuro, 10(5). doi: 10.1523/ENEURO.0087-23.2023

Bryant, K. G., Singh, B., & Barker, J. M. (2022). Reinforcement History Dependent Effects of Low Dose Ethanol on Reward Motivation in Male and Female Mice. Frontiers in Behavioral Neuroscience, 16. doi: 10.3389/FNBEH.2022.875890

Bryant, K. G., Singh, B., & Barker, J. M. (2024). Sex and individual differences in the effect of chronic low-dose ethanol on behavioral strategy selection. Alcohol, Clinical & Experimental Research, 48(1), 132–141. doi: 10.1111/acer.15218

Chen, H., He, D., & Lasek, A. W. (2015). Repeated Binge Drinking Increases Perineuronal Nets in the Insular Cortex. Alcoholism: Clinical and Experimental Research, 39(10), 1930–1938. doi: 10.1111/acer.12847

Cui, C., & Koob, G. F. (2017). Titrating Tipsy Targets: The Neurobiology of Low-Dose Alcohol. Trends in Pharmacological Sciences, 38(6), 556–568. doi: 10.1016/J.TIPS.2017.03.002

Curran-Alfaro, C. M., Bryant, K. G., Ledgister, T., Amin, S., & Barker, J. M. (2026). Sex differences in low-dose ethanol effects on motivated behavior and limbic corticostriatal activity in mice. Alcohol, Clinical & Experimental Research, 50(2). doi: 10.1111/ACER.70264

Davidson, D., Camara, P., & Swift, R. (1997). Behavioral Effects and Pharmacokinetics of Low-Dose Intravenous Alcohol in Humans. Alcoholism: Clinical and Experimental Research, 21(7), 1294–1299. doi: 10.1111/J.1530-0277.1997.TB04451.X

Daviet, R., Aydogan, G., Jagannathan, K., Spilka, N., Koellinger, P. D., Kranzler, H. R., Nave, G., & Wetherill, R. R. (2022). Associations between alcohol consumption and gray and white matter volumes in the UK Biobank. Nature Communications, 13(1). doi: 10.1038/S41467-022-28735-5

Glover, E. J., McDougle, M. J., Siegel, G. S., Jhou, T. C., & Chandler, L. J. (2016). Role for the Rostromedial Tegmental Nucleus in Signaling the Aversive Properties of Alcohol. Alcoholism, Clinical and Experimental Research, 40(8), 1651–1661. doi: 10.1111/ACER.13140

Glover, E. J., Starr, E. M., Chao, Y., Jhou, T. C., & Chandler, L. J. (2019). Inhibition of the rostromedial tegmental nucleus reverses alcohol withdrawal-induced anxiety-like behavior. Neuropsychopharmacology, 44(11), 1896–1905. doi: 10.1038/s41386-019-0406-8

Hopf, F. W., Martin, M., Chen, B. T., Bowers, M. S., Mohamedi, M. M., & Bonci, A. (2007). Withdrawal from intermittent ethanol exposure increases probability of burst firing in VTA neurons in vitro. Journal of Neurophysiology, 98(4), 2297–2310. doi: 10.1152/jn.00824.2007

Kayyal, H., Yiannakas, A., Chandran, S. K., Khamaisy, M., Sharma, V., & Rosenblum, K. (2019). Activity of insula to basolateral amygdala projecting neurons is necessary and sufficient for taste valence representation. Journal of Neuroscience, 39(47), 9369–9382. doi: 10.1523/JNEUROSCI.0752-19.2019

King, A. C., Cao, D., deWit, H., O’Connor, S. J., & Hasin, D. S. (2019). The role of alcohol response phenotypes in the risk for alcohol use disorder. BJPsych Open, 5(3). doi: 10.1192/bjo.2019.18

King, A. C., De Wit, H., McNamara, P. J., & Cao, D. (2011). Rewarding, Stimulant, and Sedative Alcohol Responses and Relationship to Future Binge Drinking. Archives of General Psychiatry, 68(4), 389–399. doi: 10.1001/ARCHGENPSYCHIATRY.2011.26

Koob, G. F., & Le Moal, M. (1997). Drug abuse: Hedonic homeostatic dysregulation. Science, 278(5335), 52–58. doi: 10.1126/SCIENCE.278.5335.52

Lea, S.E.G., Chow, P.K.Y., Leaver, L.A., McLaren, I.P.L., (2020). Behavioral flexibility: A review, a model, and some exploratory tests. Learn. Behav. 48, 173. 10.3758/S13420-020-00421-W

Li, J., Chen, P., Han, X., Zuo, W., Mei, Q., Bian, E. Y., Umeugo, J., & Ye, J. (2019). Differences between male and female rats in alcohol drinking, negative affects and neuronal activity after acute and prolonged abstinence. International Journal of Physiology, Pathophysiology and Pharmacology, 11(4), 163. Retrieved from https://pmc.ncbi.nlm.nih.gov/articles/PMC6737432/

Lin, J. Y., Arthurs, J., & Reilly, S. (2017). Conditioned taste aversions: From poisons to pain to drugs of abuse. Psychonomic Bulletin & Review, 24(2), 335. doi: 10.3758/S13423-016-1092-8

Lopez, M. F., Becker, H. C., & Chandler, L. J. (2014). Repeated episodes of chronic intermittent ethanol promote insensitivity to devaluation of the reinforcing effect of ethanol. Alcohol, 48(7), 639–645. doi: 10.1016/j.alcohol.2014.09.002

Middaugh, L. D., Kelley, B. M., Bandy, A. L. E., & McGroarty, K. K. (1999). Ethanol Consumption by C57BL/6 Mice: Influence of Gender and Procedural Variables. Alcohol, 17(3), 175–183. doi: 10.1016/S0741-8329(98)00055-X

Mitten, E. H., Souders, A., Marron Fernandez de Velasco, E., Aguado, C., Luján, R., & Wickman, K. (2025). Chronic ethanol exposure in mice evokes pre- and postsynaptic deficits in GABAergic transmission in ventral tegmental area GABA neurons. British Journal of Pharmacology, 182(1), 69–86. doi: 10.1111/bph.17335

Morikawa, H., & Morrisett, R. A. (2010). Ethanol Action on Dopaminergic Neurons in the Ventral Tegmental Area. Interaction with Intrinsic Ion Channels and Neurotransmitter Inputs. In International Review of Neurobiology (Vol. 91, Issue C, pp. 235–288). Academic Press Inc. doi: 10.1016/S0074-7742(10)91008-8

Mukherjee, A., Paladino, M. S., McSain, S. L., Gilles-Thomas, E. A., Lichte, D. D., Camadine, R. D., Willock, S., Sontate, K. V., Honeycutt, S. C., & Loney, G. C. (2023). Escalation of alcohol intake is associated with regionally decreased insular cortex activity but not changes in taste quality. Alcohol: Clinical and Experimental Research, 47(5), 868–881. doi: 10.1111/acer.15060

Przybysz, K. R., Ramirez, L. A., Pitock, J. R., Starr, E. M., Yang, H., & Glover, E. J. (2024). A translational rodent model of individual differences in sensitivity to the aversive properties of ethanol. Alcohol, Clinical and Experimental Research, 48(3), 516–529. doi: 10.1111/acer.15267

Przybysz, K. R., Shillinglaw, J. E., Wheeler, S. R., & Glover, E. J. (2024). Chronic ethanol exposure produces long-lasting, subregion-specific physiological adaptations in RMTg-projecting mPFC neurons. Neuropharmacology, 259. doi: 10.1016/j.neuropharm.2024.110098

Rabin, B. M., Hunt, W. A., & Lee, J. (1989). Attenuation and Cross-Attenuation in Taste Aversion Learning in the Rat: Studies With Ionizing Radiation, Lithium Chloride and Ethanol. In Pharmacology Biochemistry & Behavior (Vol. 31). Pergamon Press pic.

Ramirez, L. A., Przybysz, K. R., Pitock, J. R., Starr, E. M., Yang, H., & Glover, E. J. (2024). Attenuated incubation of ethanol-induced conditioned taste aversion in a model of dependence. Psychopharmacology, 241(6), 1191–1203. doi: 10.1007/S00213-024-06553-5

Riley, A. L., & Tuck, D. L. (1985). Conditioned Taste Aversions: A Behavioral Index of Toxicity. Annals of the New York Academy of Sciences, 443(1), 272–292. doi: 10.1111/J.1749-6632.1985.TB27079.X

Sheth, C., Furlong, T. M., Keefe, K. A., & Taha, S. A. (2016). Lesion of the rostromedial tegmental nucleus increases voluntary ethanol consumption and accelerates extinction of ethanol-induced conditioned taste aversion. Psychopharmacology, 233(21–22), 3737–3749. doi: 10.1007/s00213-016-4406-7

Simoes, P. M. V., Ott, S. R., & Niven, J. E. (2012). A long-latency aversive learning mechanism enables locusts to avoid odours associated with the consequences of ingesting toxic food. Journal of Experimental Biology, 215(10), 1711–1719. doi: 10.1242/JEB.068106

Taxier, L. R., Neira, S., Flanigan, M. E., Haun, H. L., Eberle, M. R., Kooyman, L. S., Markowitz, S. Y., & Kash, T. L. (2025). Retrieval of an Ethanol-Conditioned Taste Aversion Promotes GABAergic Plasticity in the Anterior Insular Cortex. Journal of Neuroscience, 45(9). doi: 10.1523/JNEUROSCI.0525-24.2024

Vetter-O’Hagen, C., Varlinskaya, E., & Spear, L. (2009). Sex Differences in Ethanol Intake and Sensitivity to Aversive Effects during Adolescence and Adulthood. Alcohol and Alcoholism (Oxford, Oxfordshire), 44(6), 547. doi: 10.1093/ALCALC/AGP048

Weafer, J., & Fillmore, M. T. (2016). Low-Dose Alcohol Effects on Measures of Inhibitory Control, Delay Discounting, and Risk-Taking. Current Addiction Reports, 3(1), 75–84. doi: 10.1007/S40429-016-0086-Y

Yang, W., Singla, R., Maheshwari, O., Fontaine, C. J., & Gil-Mohapel, J. (2022). Alcohol Use Disorder: Neurobiology and Therapeutics. Biomedicines 2022, Vol. 10, Page 1192, 10(5), 1192. doi: 10.3390/BIOMEDICINES10051192

